# Surface modification of multilayer graphene neural electrodes by local printing of platinum nanoparticles using spark ablation^†^

**DOI:** 10.1101/2023.07.30.551155

**Authors:** Nasim Bakhshaee Babaroud, Samantha J. Rice, Maria Camarena Perez, Wouter A. Serdijn, Sten Vollebregt, Vasiliki Giagka

**Author notes:** Electronic Supplementary Information (ESI) available: [details of any supplementary information available should be included here]. See DOI: 00.0000/00000000.

## Abstract

In this paper, we present the surface modification of multilayer graphene neural electrodes with platinum (Pt) nanoparticles (NPs) using spark ablation. This method yields an individually selective local printing of NPs on an electrode surface at room temperature in a dry process. NP printing is performed as a post-process step to enhance the electrochemical characteristics of graphene electrodes. The NP-printed electrode shows significant improvements in impedance, charge storage capacity (CSC), and charge injection capacity (CIC), versus the equivalent electrodes without NPs. Specifically, electrodes with 40% NP surface density demonstrate 4.5 times lower impedance, 15 times higher CSC, and 4 times better CIC. Electrochemical stability, assessed via continuous cyclic voltammetry (CV) and voltage transient (VT) tests, indicated minimal deviations from the initial performance, while mechanical stability, assessed via ultrasonic vibration, is also improved after the NP printing. Importantly, NP surface densities up to 40% maintain the electrode optical transparency required for compatibility with optical imaging and optogenetics. These results demonstrate selective NP deposition and local modification of electrochemical properties in neural electrodes for the first time, enabling the cohabitation of graphene electrodes with different electrochemical and optical characteristics on the same substrate.

## 1 Introduction

In recent years, the combination of complementary methods such as optical and electrical neural recording and stimulation has enabled a deeper understanding of the brain and deciphering neural behavior with the goal of advancing the treatments and therapies for disorders and diseases related to the nervous system.

A combination of various modalities can be used to monitor the neural response of multiple neurons. Optical imaging (calcium imaging and two-photon imaging) ^1,2^ together with electrophysiology, the method used for neural activity recording, have been employed to target specific biological structures and identify cell types. The combination of optogenetics with electrophysiology has also attracted great attention in neuroscientific research in recent years to pave the way towards a much deeper understanding of the nervous system^2–4^.

However, conventional metal-based electrodes, mostly used in neural interface devices, are not the best candidates to combine electrical and optical neural measurement methods. One challenge of using such electrodes for the combination of optical imaging and electrophysiology is the obstructed field of view due to metal opacity. Another challenge of using such electrodes for the combination of electrical recording and optical stimulation is that photo-induced artifacts might be generated due to photoelectrochemical effects which might interfere with the recorded electrical signal ^5,6^. Therefore, the development of transparent conductive materials has increased rapidly to substitute metal electrodes.

Graphene as a potential electrode material has a high thermal/electrical conductivity and broad-spectrum transparency^7^. In addition, using a chemical vapor deposition (CVD) technique for graphene growth makes it compatible with micro-fabrication process steps.

Monolayer CVD graphene with high optical transparency has shown compatibility with optical imaging and optogenetics^8^. However, undoped monolayer graphene suffers from low sheet conductivity^9^ and a low charge storage capacity (CSC) due to the dominance of its small quantum capacitance^10^. Multilayer CVD graphene electrodes have been recently reported using a transfer-free fabrication process^11,12^. Although using multiple layers of graphene increases the quantum capacitance and consequently reduces the impedance and leads to higher CSC^11^, increasing the number of layers has only impact on the quantum capacitance up to a threshold of 6 layers ^10^, while each added layer reduces the graphene’s optical transparency^11,13^.

Moreover, scaling down the electrode size is necessary to selectively record signals from targeted neurons^14,15^. However, a size reduction is accompanied by an increase in impedance, which causes an increase in the amount of noise (voltage) from the electrodes:

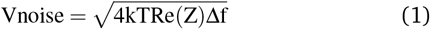

where k is Boltzmann’s constant, T is the temperature, Z is the electrode impedance, and Δf is the frequency band of interest^16,17^.

Furthermore, high-impedance electrodes require higher voltages to drive the desired stimulation currents to trigger a neural response, making their use impractical in implantable systems operating with a limited power budget.

Thus, to obtain a low impedance with small-size electrodes, various strategies have been investigated. The surface modification techniques mostly used in the literature rely on either materials that increase the surface roughness (topographical approach) or on materials that improve the impedance and CSC by additional electrochemical means (chemical approach)^18^.

Various materials have been applied to modify the graphene electrode surface, including nanoparticles (NPs)^19^, carbon nanotubes (CNTs)^20,21^, and conductive polymers such as poly(3,4-ethylene dioxythiophene) (PEDOT)^22,23^. NPs, specifically, have a larger surface-to-volume ratio compared to bulk materials. Therefore, the deposition of NPs on electrodes has been shown to lower the impedance^24^. Furthermore, the surface topography of electrodes can improve the cell adhesion to the electrode surface and, therefore, improve signal quality^25^.

Gold (Au) NPs are highly conductive and therefore widely used in the field of biosensors, multielectrode arrays, and microelectrodes. Graphene with Au NPs has been fabricated by doping the graphene surface with AuCl_3_^26^. This method showed a greater reduction in sheet resistance than doping with HNO_3_, while the achieved optical transmittances were similar.

Electrodeposition of platinum (Pt) NPs on reduced graphene oxide (rGO)^27,28^, and on functionalized graphene sheet^29^ has been shown to enhance the electroactivity in fuel cell and biochemical sensor applications. Recently, graphene neural electrodes with electrodeposited Pt NPs showed a reduction in the impedance and a CV enlargement for an increased Pt deposition time^19^. It has been suggested that creating an alternative conduction path with Pt NPs at the electrode-electrolyte interface could increase the small quantum capacitance^19^. Moreover, Pt is a material conventionally used for stimulation electrodes due to its high charge injection capability. Therefore, adding Pt NPs on the graphene surface was expected to increase the amount of faradaic charge transfer over the electrode-electrolyte interface and thus increase the CSC^19^.

The current techniques used for surface modification of the electrodes with Pt NPs are mostly based on chemical or electrochemical depositions. The NP coating formation by these methods is highly time-consuming and hard to control, which results in limited reproducibility and mass production^30^. Moreover, the NP coating process step optimised and integrated into the fabrication process of specific electrodes with specific sizes and materials may not easily be applicable to other types of electrodes^18^. Also, these chemical process steps might result in a lower purity as they tend to leave residues on the electrodes. Furthermore, single (individual) electrode surface modification is not always possible and usually comes at a high cost and process complexity.

The aim of this paper is the direct surface modification of multilayer graphene electrodes with Pt NPs using the spark ablation method to enhance the electrochemical performance of the electrode. This versatile method is based on gas-phase electro-deposition, which prevents the exposure of the electrodes to any chemicals. It is capable of single-step local NP impaction printing and is compatible with the existing microfabrication process as a post-processing step. Due to its local nature, this technique opens up new possibilities in neural interface design for multimodal tissue interaction. For instance, it enables the coexistence of smaller size electrodes, which need better electrochemical characteristics, with larger electrodes, requiring higher optical transparency, on the same substrate.

## 2 Methods

### 2.1 Sample preparation

#### 2.1.1 Fabrication process

The fabrication process of the multilayer graphene electrode, as shown in Fig. 1(a), starts with 300 nm thermal wet oxide growth on the front side of a silicon (Si) wafer at 1000 °C. Next, 50 nm molybdenum (Mo) is sputter-deposited at 50 °C on the oxide layer. The Mo layer is then patterned and etched to serve as the catalyst layer for the following graphene growth. Mo etch is performed using an inductively coupled plasma (ICP) etcher with 50 W RF power, 500 W ICP power, 5 mTorr pressure, 25 °C temperature, and 30 and 5 sccm Cl_2_ and O_2_ gas flows, respectively. Then, graphene is selectively grown on pre-patterned Mo using a chemical vapor deposition (CVD) process (using an Aixtron Black Magic Pro tool) at temperatures of 935 °C, 25 mbar pressure, and using 960, 40, and 25 sccm of Ar, H_2_, and CH_4_ gas flows, respectively, and cooled to room temperature under an Ar atmosphere. The growth time is 20 minutes which results in 7 graphene layers as shown in^11^.

**Fig. 1.**
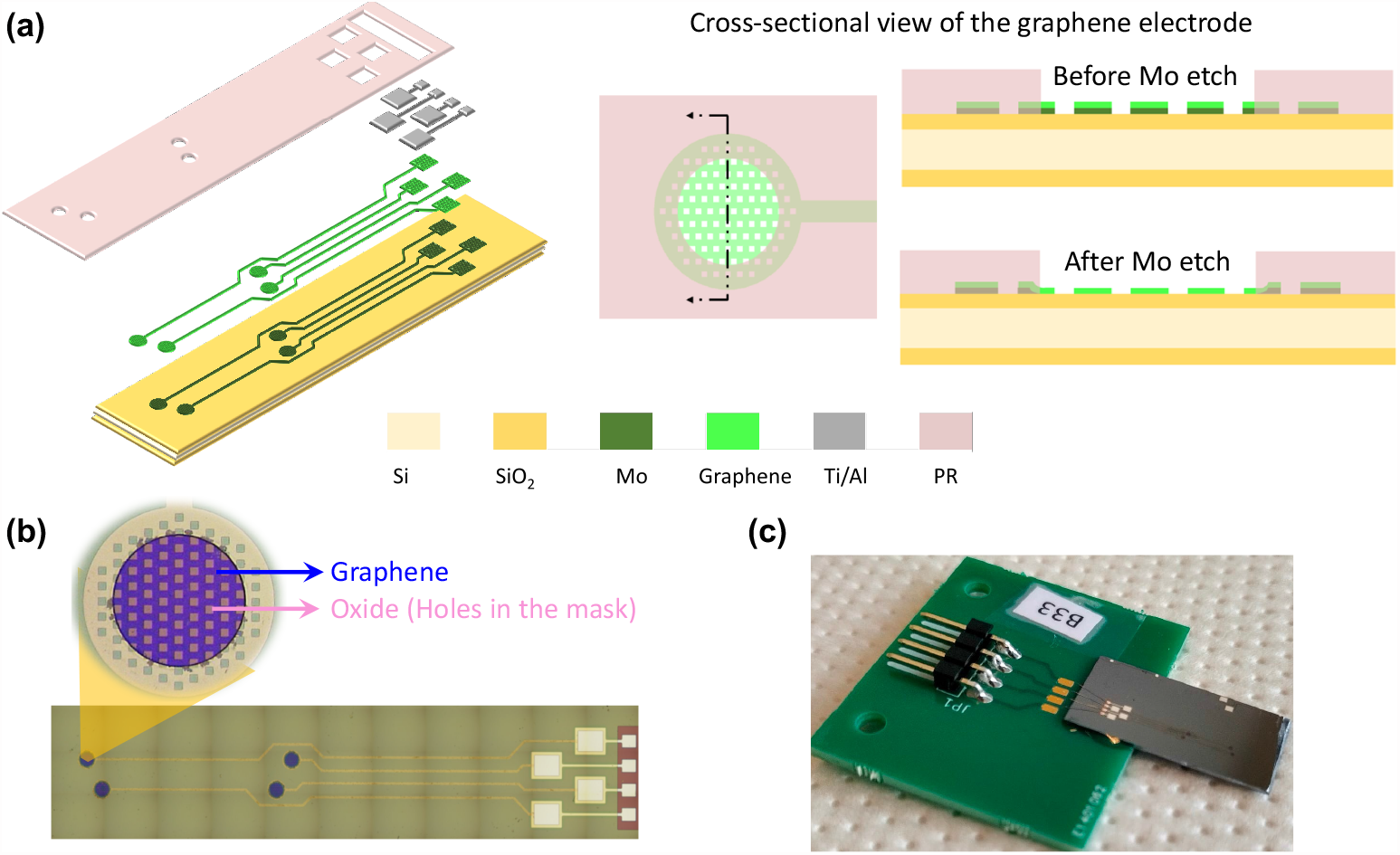
(a) Fabrication process steps of graphene-based neural electrodes on a Si substrate. First, the oxide is deposited on the front side of a Si wafer followed by Mo deposition and patterning. Then, graphene is grown on a Mo catalyst. Next, Al/Ti is deposited and patterned on the contact pads. Finally, PR is spin-coated and patterned as an insulation layer and Mo is removed from the electrode sites, leaving graphene contacts (as shown in the cross-sectional view of the graphene electrode). (b) Optical image of the final device with four electrodes. (c) The final device attached and wire-bonded to a PCB.

Then, an aluminum (Al) / titanium (Ti) stack is sputtered and photolithographically patterned to allow for wire bonding on the contact pads. The Al/Ti layer is then etched in a 0.55% concentration of hydrofluoric acid (HF) to remove this layer from the whole wafer except the contact pads. Next, the photoresist (PR) is spin-coated as an insulation layer and patterned on the electrodes and contact pads. Finally, Mo underneath the graphene electrodes is etched using wet etching in hydrogen peroxide (H_2_O_2_), leaving the graphene in the exact same location as defined by the catalyst. This is shown in the cross-sectional view of the graphene electrode before and after etching Mo in 1(a). The optical image of the final device with four electrodes is shown in Fig. 1(b). The electrode diameter is 340 μm which leads to 68320 *μ*m^2^ surface area after subtracting the holes’ surface area (these holes are considered in the mask design as explained in^11^).

At the end of the fabrication process, the Si wafer is diced and an epoxy die adhesive is used to attach one device to a printed circuit board (PCB) for further testing (Fig. 1(c)). Next, the Al contact pads are Au wire-bonded to the PCB pads. The contact pad and the attached wire are both covered with a drop of Polydimethylsiloxane (PDMS) for mechanical protection. At this stage, some preliminary measurements are performed to characterize the graphene electrodes (pre-NP measurements). Next, the electrodes are ready for NP printing and post-NP measurements.

#### 2.1.2 NP printing

A spark ablation method is employed to print NPs on the electrodes. The process consists of three steps: generation, particle processing, and deposition as illustrated in Fig. 2. A generator (VSP-G1) is connected to a prototype nanostructured material printer (VSP-P0) (VSPARTICLE BV, the Netherlands). The generator initiates periodic electrical discharges between two metal rod electrodes of a desired conductive material and an inert gas flow (Nitrogen (N_2_)) carries the NPs to the substrate in the deposition chamber.

**Fig. 2.**
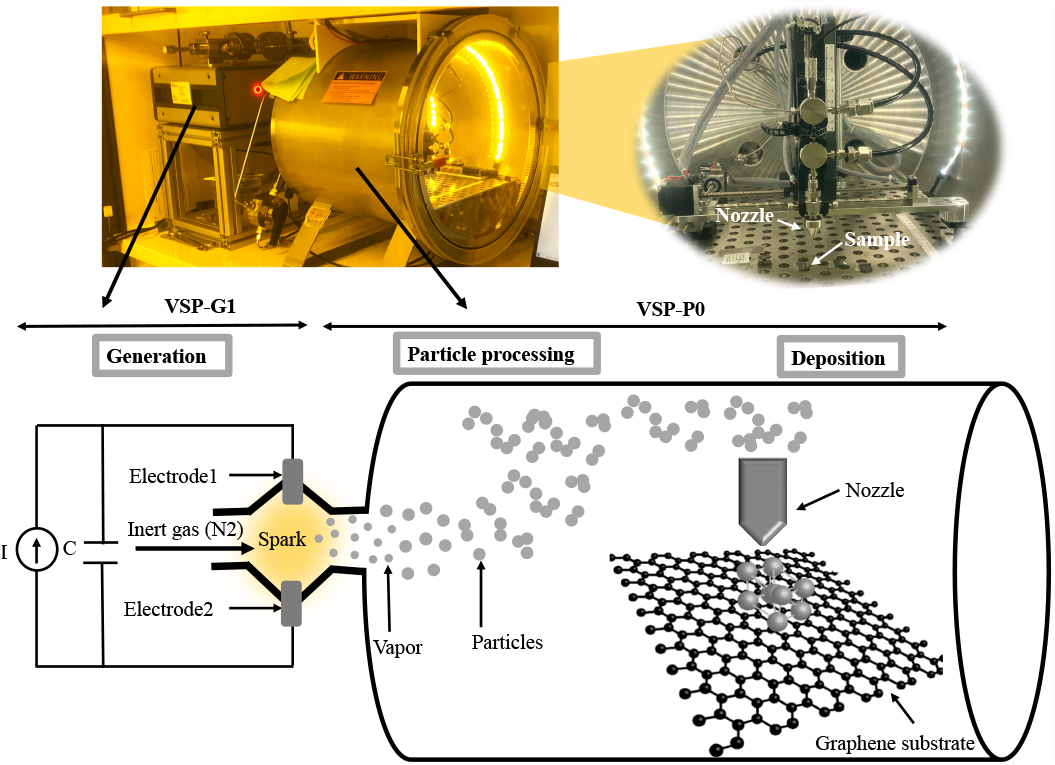
Schematic view of the spark ablation method system (VSPARTI-CLE BV, the Netherlands) used for NP printing.

The substrate is mounted on the stage in the vacuum chamber perpendicular to a 3D-printed converging nozzle with a 0.35 mm diameter as shown in Fig. 2. This nozzle is connected to motors that can navigate in the x,y, and z directions, creating a local printing process. The nozzle aerodynamically focuses the NPs and deposits them on the substrate by impaction^31^. In this work, Pt (99.9% purity) rod electrodes with diameters of 3 mm are used to create Pt NP coating on graphene electrodes.

#### 2.1.3 Optimisation of NP printing settings

The printer settings are optimised to get three different surface densities of NPs, namely 15%, 30%, and 40%. The spark is generated with 1 kV and 3 mA current and the printings are performed under ambient pressure with nitrogen (99.995% purity) as a carrier gas at a flow rate of 1.5 l/min.

First, NP printing is performed on silicon dies from a nozzle height of 0.5 mm in single-line patterns by varying printing speeds. Then, the Si dies are inspected by scanning electron microscopy (SEM), and the corresponding images are analyzed to calculate the obtained NP densities per each printing speed. Finally, the printing speed to achieve the required NP density is chosen.

For each printer setting, the resulting NP density is determined by averaging the surface density of three SEM images from the same deposition batch. These images are taken with a 2 kV electron beam and 50000x magnification (using a Hitachi Regulus 8230). First, the grey-scale SEM images are converted to black and white (binary) images using MATLAB R2019b (of MathWorks). The pixels whose value is above a certain threshold are replaced with white, representing the area covered with NPs, and the pixels with values below the threshold are considered black. Otsu’s method^32^ is used to determine the optimal threshold to convert the grey-scale images into binary images. The percentage of white pixels out of all pixels is considered the density of the NPs.

To ensure that the width of the printed NP line is sufficient to cover the electrode surface, it is necessary to measure the line width. SEM images taken at low magnification (x50) at a 2kV accelerating voltage are used for this purpose. The grey-scale images are converted to binary images through Otsu’s thresholding method. The data matrix includes columns of 0s and 1s and the longest series of 1s across a column is considered the width. The average length of that data matrix is then used as the width of the printed line. This value is subsequently converted to millimeters using ImageJ (an image analysis program developed at the National Institute of Health^33^).

It should be noted that the printed line should not conduct electricity to prevent enlarging the circular electrode surface area into an extended line. To ensure low conductivity of the printed NP lines, a conductivity measurement is performed using a four-point probe (Cascade Microtech probe station) on gold transfer length method (TLM) structures covered with an NP line with the same density printed over them.

### 2.2 Electrode characterization

#### 2.2.1 Raman spectroscopy

Raman spectroscopy is used to investigate the effect of NP printing on the graphene lattice structure. For the Raman characterization, a Renishaw inVia system with a red HeNe laser of 633 nm is used. Raman spectroscopy is performed before and after NP printing on the electrode surface.

#### 2.2.2 Surface roughness

The NP surface density can only indicate the 2D distribution of the NPs. To indicate a 3D NP distribution over the electrode surface, surface roughness measurements are performed. A high surface roughness has been shown to increase the CSC due to an increase in the electrochemical surface area of the electrode^34^. The surface topography of the Pt NPs printed with multiple surface densities is investigated through atomic force microscopy (AFM; Ntegra). Samples are prepared by printing Pt NP lines on a silicon die. The topography of these samples is tested at a frequency of 0.50 Hz in semi-contact mode with a scan size of 10 *μ*m by 10 *μ*m. This measurement is performed 5 times for each sample at different locations throughout the printed NP lines but as far as possible from the line edges. The AFM data is further processed using Gwyddion applying polynomial correction of the background^35^.

#### 2.2.3 Optical transmittance measurements

The optical transparency of different NP surface densities is assessed for wavelengths in the range of 300 nm to 900 nm (using a Perkin Elmer Lambda 950 UV/Vis spectrophotometer, Waltham, Massachusetts). NPs are printed directly on glass slides (1 sample per surface density). To create a sufficiently large area covered with NPs for this measurement, the printer’s nozzle follows a laddered path, i.e., a line is printed along the x-direction followed by a step in the y-direction. Since the line width for each NP density is determined through the method discussed previously, a logical step at the y-direction is considered for each NP density to minimize the chance of overlapping with the previous line.

#### 2.2.4 Electrochemical impedance spectroscopy

Electrochemical impedance spectroscopy (EIS) is employed to assess the electrochemical properties of the graphene electrodes with and without NP coating. The measurements are performed in a phosphate-buffered saline (PBS) solution in a three-electrode configuration set-up using a potentiostat (Autolab PG-STAT302N). A Pt electrode (3 mm diameter (BASI Inc.)) is used as a counter electrode, a leakless miniature silver/silver chloride (Ag/AgCl) (eDAQ) as a reference electrode, and the graphene electrodes(with and without NPs) fabricated in this work as the working electrodes. A 10-mV RMS sinusoidal voltage is applied between the working and reference electrodes and the current between the working and counter electrodes is measured^36^. Finally, the impedance magnitude and phase angles are plotted as a function of frequency ranging from 1 Hz to 100 kHz.

#### 2.2.5 Cyclic voltammetry

Cyclic Voltammetry (CV) is an electrochemical surface analysis technique used for investigating charge transfer reactions of an electrode surface. CSC is calculated as the time integral of the CV curve and is reported as charge per electrode surface area. This value estimates the total charge transferred to the electrode and has been used as a common practice to compare stimulation electrodes^36,37^. CV measurement is performed using the same three-electrode setup. To ensure a safe measurement for both materials (Pt and graphene) and facilitate the comparison between pre-NP and post-NP measurements, the overlap (−0.6 V to 0.6 V) between the previously used water window ranges for graphene (−0.8 V to 0.6 V) and Pt (−0.6 V to 0.8 V) is chosen^11^. The measurements are performed at various scan rates (0.1 V/s, 0.2 V/s, 0.6 V/s, and 1 V/s) 3 times to ensure that the third stabilized cycle is used for the calculation of the CSC. Both the total and cathodic CSC are calculated and expressed in *μ*C/cm^2^ after dividing the calculated charge (based on the third scan) by the electrode surface area.

#### 2.2.6 Voltage-transient measurements

Voltage-transient (VT) measurements are used to estimate the maximum charge that can be injected by an electrode by applying a constant-current stimulation pulse^34,36,37^. This measurement is performed in the same three-electrode setup as well. A cathodic-first biphasic symmetric current pulse (1 ms pulse width, 100 *μ*s interphase delay) is applied between the working and counter electrodes in the PBS solution. The voltage between the working electrode and the reference electrode is then evaluated. This voltage consists of a resistive voltage drop at the beginning of the cathodic pulse followed by a gradual voltage decrease due to the capacitive charging of the electrode-electrolyte interface. The interface polarization of the electrode is evaluated by eliminating the resistive voltage drop from the minimum voltage. The interface polarization should not exceed the water window extracted from CV. The maximum cathodic-current amplitude is the maximum current in which the interface polarization reaches the cathodic water window. Finally, the maximum charge-injection capacity (CIC) of the electrode is calculated based on the maximum current amplitude multiplied by the pulse width and divided by the electrode surface area^34^.

### 2.3 Stability assessments

Printing NPs on the graphene electrode surfaces is performed to reduce the electrode impedance and increase the CSC. This electrochemical improvement should remain stable for the lifetime of the device. Coating stability includes both electrochemical and mechanical stability, meaning the electrode should maintain its electrochemical improvement and the coating should not delaminate from the electrode surface. Therefore, the electrochemical and mechanical stability of the NPs must be verified prior to any electrode implantation in the body.

To this aim, continuous CV and VT tests are performed to ensure the electrochemical stability of the electrodes. Finally, an ultrasonic vibration test is performed to evaluate the mechanical stability of the NP coating.

#### 2.3.1 Continuous CV measurement

Continuous CV measurements are commonly used to evaluate the electrochemical stability of the electrodes^38^. Three samples per each electrode type are subjected to 500 CV scanning cycles at a fast scan rate of 1 V/s. EIS measurements are then performed to evaluate any changes in the impedance. The impedance of the electrodes at 1 kHz is reported before and after this test. To additionally investigate whether the Pt NPs are present on the electrode surface after 500 CV cycles, energy-dispersive X-ray spectroscopy (EDX) measurement is performed on the electrode with NP coatings.

#### 2.3.2 Continuous VT measurement

A continuous VT test is performed by applying cathodic-first biphasic current pulses to the electrode. A current amplitude of 2.5 *μ*A with a 1 ms pulse width and 1 *μ*s interphase delay with a frequency of 333 Hz is applied. The number of cycles is kept at 500,000 as the Mo layer underneath the graphene tracks started to corrode. The test is conducted for two electrodes: one graphene electrode without any NP coating and one graphene electrode with 40% NP coating. The characterization of the electrodes is performed before and after the continuous VT test by performing Raman spectroscopy, EIS, and CV measurements.

#### 2.3.3 Ultrasonic test

The stability of the NPs on the electrode surface is additionally tested by ultrasonic vibration using a digital ultrasonic cleaner (HBM Machines). Four electrodes, two graphene electrodes without any coatings, and two graphene electrodes with 40% Pt NPs are submerged in a water bath of 250 ml at 30 W power, 22 kHz frequency for 2 minutes. Optical images of the electrode surface are taken before and after this test. EIS measurements are also performed as any changes in the impedance might reveal a change in the surface properties and the delamination of NPs from the electrode surface.

## 3 Results

### 3.1 NP printing

Fig. 3(a) shows the SEM images of NPs printed on Si dies with the required surface densities. The corresponding binary image of 40% Pt NPs is also depicted. The settings and parameters used for Pt NP printing are shown in Table S1 in the supplementary notes. The resulting printed line width, calculated from the binary images for the various NP densities, is also reported in this table. In all cases, the lines are wide enough to ensure that NPs are printed on the entire electrode surface with a diameter of 340 *μ*m.

**Fig. 3.**
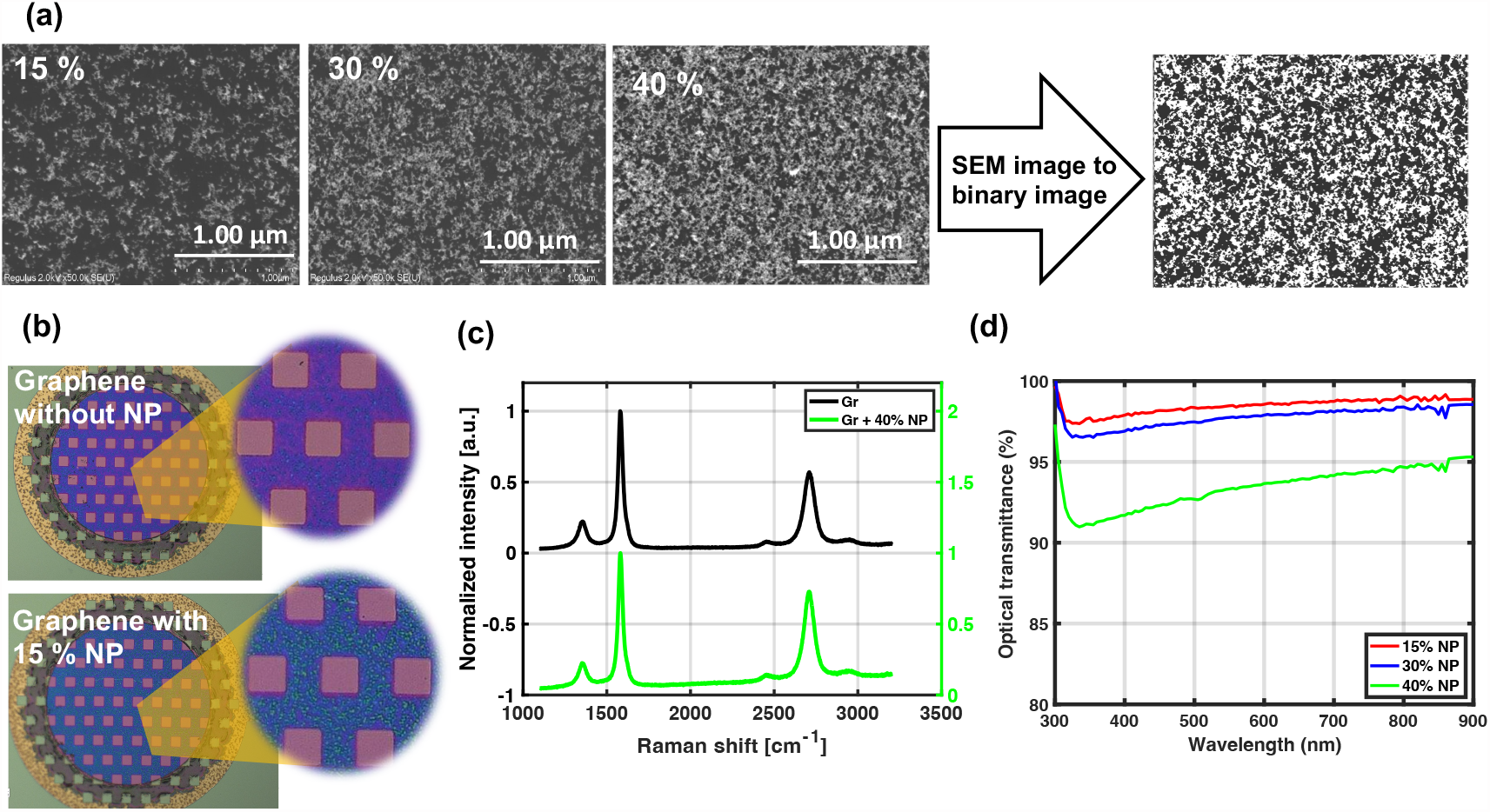
(a) The original grey-scale SEM images after calculating the surface density of NPs, together with the corresponding binary image of 40% NP density, (b) The optical image of graphene electrode before and after 15% NP printing, (c) Raman spectroscopy measurement of graphene without and with 40% NPs, (d) Optical transmittance of different NP densities (15, 30, and 40%) on glass after removing the effect of the glass slide.

Furthermore, the results of the conductivity measurement can be found in Fig. S1 in the supplementary notes. The measurement on the printed Pt NP lines over gold TLM structures shows a significant current flow for NP surface densities of 50%. Therefore, the NP surface density is kept below 40% for this study to prevent any potential extension of the graphene electrode surface to the printed line, as this would influence the electrochemical tests. Conductivity measurements for these lower densities, discussed in the supplementary notes, indicated this has not been the case.

Fig. 3(b) shows the optical images of a graphene electrode before and after 15% Pt NP printing. NPs can be seen in the zoomed-in image and an obvious color change in the electrode surface is observed as a result of the presence of NPs.

### 3.2 Electrode characterization

#### 3.2.1 Raman Spectroscopy

The Raman spectrum of the graphene with and without Pt NPs is displayed in Fig. 3(c). The spectrum of graphene with 40% Pt NPs is shown in green and the spectrum before printing NPs on graphene is displayed in black as a reference. Three distinct peaks can be observed for both spectra: a D peak at 1354 cm^−1^, a G peak at 1582 cm^−1^ related to the sp2 C-C bonds forming the graphene lattice, and a 2D peak around 2709 cm^−1^. No differences are observed in the average intensity ratio of the D to G peaks (I(D)/I(G)) after NP printing (0.19 for graphene and 0.17 for graphene with Pt NPs), implying that Pt NPs did not affect the defect density of graphene. A possible explanation is that nanoparticles are small and sparsely distributed over the surface and they do not have sufficient kinetic energy to cause defects.

The mechanism of printing NPs probably results in physically adsorbed clusters causing NPs to have minimal interaction with the graphene lattice.

#### 3.2.2 Surface roughness

The surface roughness measurement on the printed Pt NPs on Si dies is performed for five samples per surface density. The average and standard deviation of RMS and mean surface roughness for each NP surface density are reported in Table S2 in the supplementary notes. The reported values show a higher average for both RMS and mean roughness by increasing the NP surface density from 15 to 40% (from 9 to 14.67 nm for RMS and from 7.1 to 12 nm for mean surface roughness). An increase in the standard deviation of both RMS and mean surface roughness by increasing the NP density confirms the non-uniformity of printed NPs over the surface. It should be noted that the surface roughness of multilayer graphene electrodes with the same growth condition was reported to be 6.75 nm^11^ which is smaller than the surface roughness of printed Pt NPs.

#### 3.2.3 Optical transmittance measurements

The optical transmission of different NP surface densities versus the wavelength is shown in Fig. 3(d). The optical transmittance measurement was not directly performed for 40% NP surface density. The result shown in the graph is the interpolation of the optical transmittance of 30% and 50% (not shown) NP densities. As shown in the graph, the optical transmittance of the NPs with 40% surface density is still above 92% over a wide range of wavelengths.

#### 3.2.4 Electrode impedance spectroscopy

EIS measurements are performed for graphene electrodes without and with NP coatings with various surface densities. The average impedance magnitude and phase angle of 7 graphene electrodes without NPs and 5 electrodes with NPs per each NP surface density are shown in Fig. 4(a). A 2-times reduction in the impedance magnitude at 1 kHz is observed after adding the NPs with 15% surface density to the graphene-electrode surface. The graphene-electrode impedance decreased even further by printing 40% NPs from 31.45 kΩ to 7.26 kΩ (leading to a 4.5 times impedance reduction). The average impedance and the area-normalized impedance of the electrodes at 1 kHz before and after NP printing for various NP surface densities are represented in Table 1.

**Table 1.** Impedance, CSC, and CIC of graphene, without and with Pt NPs with surface densities of 15%, 30%, and 40%.

| Measurement results |  |  | Graphene | Graphene + 15% Pt NPs | Graphene + 30% Pt NPs | Graphene + 40% Pt NPs |
| --- | --- | --- | --- | --- | --- | --- |
| Impedance at 1 kHz (kΩ) |  |  | 31.45 ± 5.88 | 14.29 ± 0.95 | 9.32 ± 0.80 | 7.26 ± 0.90 |
| Area-normalized impedance (Ω.cm²) |  |  | 21.49 ± 4 | 9.8 ± 0.6 | 6.4 ± 0.5 | 4.96 ± 0.6 |
| CSC (μC/cm²) | Total | 1V/s | 248 ± 185 | 551 ± 73 | 745 ± 91 | 954 ± 103 |
|  |  | 0.6V/s | 349 ± 272 | 778 ± 109 | 959 ± 159 | 1199 ± 99 |
|  |  | 0.2V/s | 817 ± 666 | 1887 ± 268 | 1960 ± 504 | 2444 ± 368 |
|  |  | 0.1V/s | 1566 ± 1277 | 3497 ± 495 | 3399 ± 1062 | 4365 ± 821 |
|  | Cathodic | 1V/s | 78 ± 52 | 345 ± 84 | 469 ± 73 | 651 ± 45 |
|  |  | 0.6V/s | 95 ± 63 | 496 ± 146 | 625 ± 118 | 878 ± 88 |
|  |  | 0.2V/s | 159 ± 102 | 1211 ± 439 | 1327 ± 367 | 1993 ± 429 |
|  |  | 0.1V/s | 233 ± 146 | 2197 ± 849 | 2207 ± 746 | 3614 ± 892 |
| Max. current amplitude (μA) |  |  | 5.7 ± 1.5 | 8.6 ± 1.3 | 12.5 ± 1.5 | 20.9 ± 1.2 |
| CIC (μC/cm²) |  |  | 8.4 ± 2.1 | 12.6 ± 1.9 | 18.3 ± 2.2 | 30.6 ± 1.8 |

**Fig. 4.**
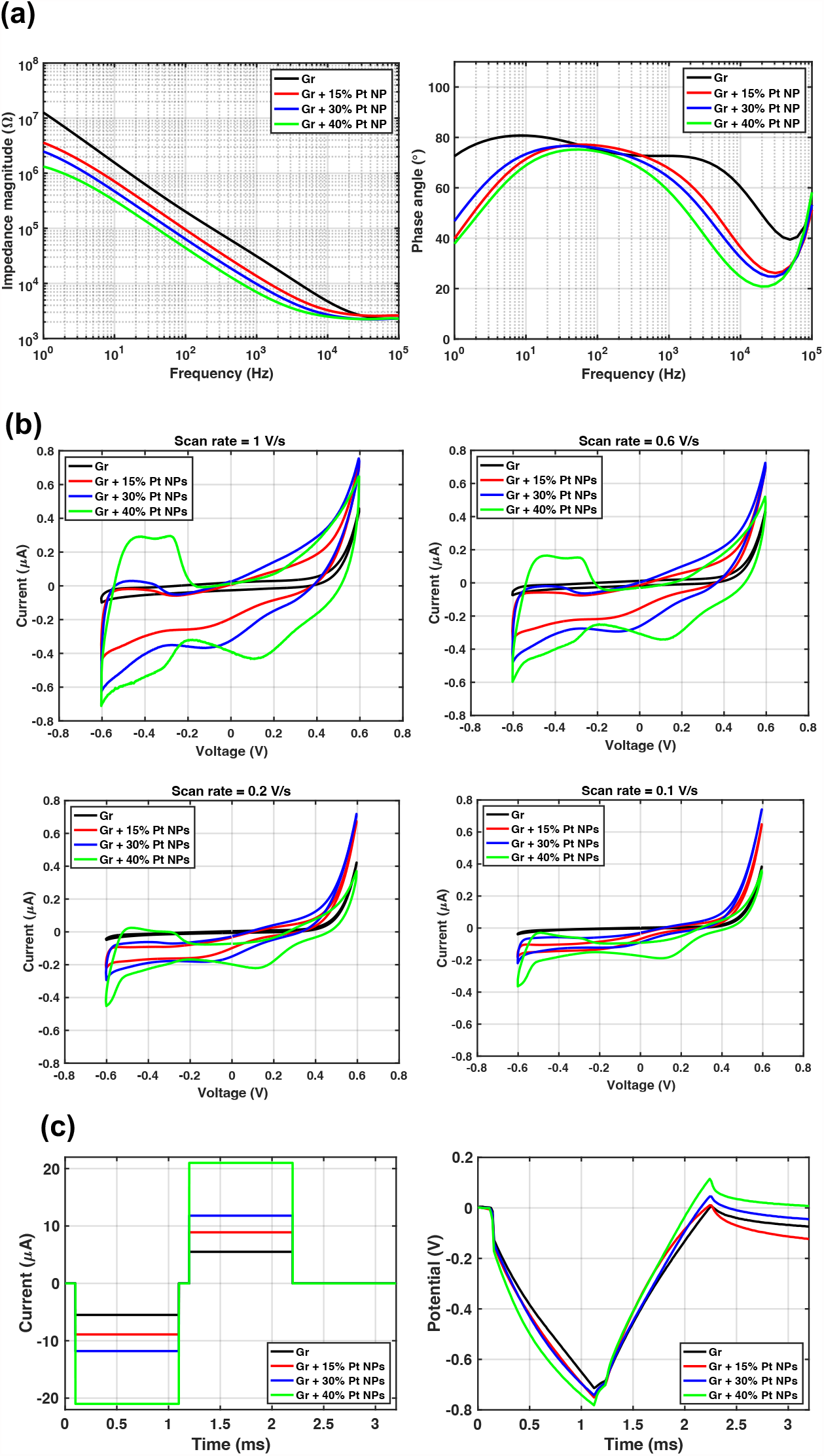
(a) Average EIS results of 7 samples per graphene without and 5 samples with NP with various surface densities, (b) CV curves of the median sample of each category of graphene without and with NPs with different surface densities, (c) The median of the maximum-amplitude current pulse applied to 5 electrodes per each category of different NP densities with the corresponding VT measurement.

#### 3.2.5 Cyclic voltammetry

The CV curves of the median sample of each sample group including graphene electrodes with and without NP coatings for various scan rates are shown in Fig. 4(b). An enlargement of the CV curve is clearly observed after increasing the surface density of the NPs. In addition, oxide reduction peaks at 0.1 V and hydrogen adsorption peaks at − 0.4 V indicate the engagement of Pt NPs in the charge-transfer process at the electrode/electrolyte interface^37^. The calculated total and cathodic CSC of each electrode are represented in Table 1. As shown, a CSC increase correlates with an increase in the surface density of NPs on the electrode surface, and a 40% NP coating improves the CSC 15 times compared to that without NPs (from 233 *μ*C/cm^2^ to 3614 *μ*C/cm^2^).

#### 3.2.6 Voltage-transient measurements

The maximum current amplitudes and measured voltages for the median samples of each sample group with respect to the (Ag/AgCl) reference electrode are shown in Fig. 4(c). An increase is observed in the maximum current pulse amplitude applied to the electrode by increasing the NP surface density, consequently resulting in an increase in the calculated maximum CIC of up to 3.5 times, as shown in Table 1.

### 3.3 Stability assessment

#### 3.3.1 Continuous CV measurement

Continuous CV tests are performed for graphene electrodes with and without NPs for three electrodes per each type for 500 cycles. The CV curves of one representative graphene electrode without NP and one with 40% Pt NP after 3, 250, and 500 CV cycles are shown in Fig. 5(a). A slight increase in the area of the CV curve (4.7% increase in total CSC) of graphene electrodes without coating is observed. The corresponding EIS measurement before and after 500 cycles of CV are also shown for these electrodes in Fig. 5(a). A small reduction (only 3%) in the impedance of the graphene electrode is observed which might be related to surface cleaning and contamination removal of the electrode surface.

**Fig. 5.**
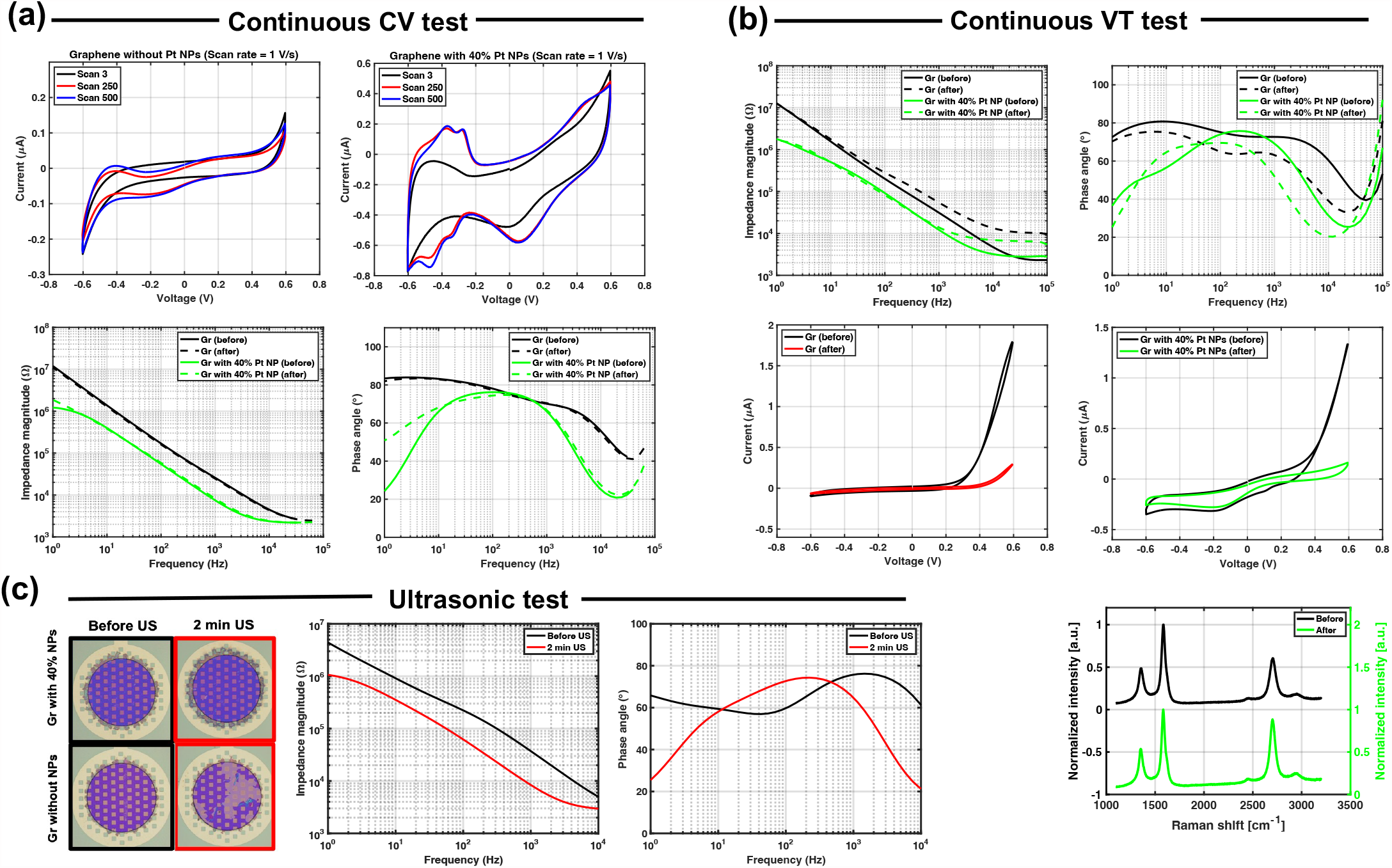
Stability assessment results. (a) Continuous-CV test results: CV curves of graphene without NP and graphene with 40% NP after 3, 250, and 500 cycles, and impedance magnitude and phase plot of graphene without NP and with 40% NP before and after 500 cycles of CV, (b) Continuous VT test results: impedance magnitude and phase plot of graphene electrodes without and with 40% NP coatings before and after 500,000 cycles of VT test, CV curves of graphene electrodes without and with 40% Pt NP coatings before and after continuous VT test, and Raman spectra of a graphene electrode with 40% NP coatings before and after continuous VT test, (c) Ultrasonic test results: an optical image of graphene electrodes with and without NPs before and after 2 minutes of ultrasonication, and impedance magnitude and phase plot of graphene electrode with 40% NP density before and after 2 minutes of ultrasonication.

A CV enlargement is observed for 7 out of 9 graphene electrodes with Pt NP coating after 500 cycles of CV (shown in Table S3 in the supplementary notes). A shoulder at 0.25 V is observed that corresponds to oxide formation, and a peak at 0.10 V is attributed to the oxide reduction. Peaks for hydrogen adsorption and desorption were also observed around -0.40 V as reported previously^37^. Comparing the CV curves after 250 and 500 cycles shows that the CV curve seems to be stabilized and no significant change is observed after 250 cycles. Although the CV enlargement is observed for graphene electrodes with various Pt NP densities after 500 CV cycles, the electrodes with 40% NPs show the largest increase in CSC of around 16.9%. This can be related to more oxidation and reduction as a result of more Pt NPs and, therefore, higher peaks in CV. The CV curve reduction of 2 out of 9 electrodes is inconclusive and cannot be attributed to the delamination of Pt NPs. To draw any conclusions, more samples should be tested and the number of CV cycles should be also increased.

The impedance of 7 out of 9 graphene electrodes with Pt NPs shows an average of 15% increase. This increase could be attributed to Pt NP delamination which could increase the impedance. However, if there is any delamination of NPs the CSC is expected to decrease. It might be argued that the appearance of the peaks related to oxidation and reduction compensates for the NP delamination. However, if this were the case, the delamination is expected to continue even after 250 CV cycles.

Finally, the EDX measurement on the electrode surface after a continuous CV test (shown in Fig. S2 in the supplementary notes) confirms the presence of the coating by showing Pt peaks corresponding to Pt NPs.

#### 3.3.2 Continuous-VT measurement

EIS plots of graphene without and with 40% Pt NPs are shown in Fig. 5(b) before and after this test. An increase in the impedance at 1kHz (from 34.53 kΩ to 54.88 kΩ) is observed for the graphene electrode without NP coating which could be related to the small delamination of graphene from underlying oxide. This delamination could be responsible for the slight increase also observed in the impedance at 1 kHz for graphene with 40% NP coatings (from 6.7 kΩ to 9.75 kΩ). A significant delamination of NP from the graphene layer is unlikely to be occurring, as the impedance curve remains about an order of magnitude lower for the NP-coated electrodes throughout the frequency spectrum. An impedance increase at high frequencies could be attributed to the corrosion of Mo underneath the graphene tracks.

CV curves for graphene electrodes without and with 40% NP coating are shown in Fig. 5(b). In both cases a reduction in the CV area is observed after 500,000 cycles of the VT test, indicating some electrochemical change in the graphene electrode. However, the CSC for the NP-coated electrodes remains higher than the one of graphene only, further suggesting the presence of NP still after the continuous-VT test.

The Raman spectra acquired on the graphene electrode with 40% NP coating before and after this test show three graphene characteristics, confirming the presence of graphene after the continuous-VT test as shown in Fig. 5(b). However, the I(D)/I(G) ratio slightly increased from 0.2 to 0.35 probably due to the defects induced into the graphene lattice. This ratio did not change for the graphene electrode without any coatings.

#### 3.3.3 Ultrasonic test

Some of the samples are additionally subjected to ultrasonic treatment to investigate its effect on the NP adhesion to the electrode surface. However, the test samples are not optimized for this test, hence the test could not be performed for long durations due to the delamination of graphene from the underlying oxide layer on the test samples (shown in Fig. 5(c)). Nevertheless, the samples with NPs remain stable after 2 minutes of ultrasonic treatment as shown in Fig. 5(c). The impedance magnitude of the graphene sample with NPs (Fig. 5 (c)) shows a decrease after 2 minutes of ultrasonication. Graphene delamination from the underlying oxide starts after 2 mins, however, for NP-coated electrodes which presented higher mechanical stability, the treatment was continued. EIS measurements after 10 minutes of ultrasonication indicate that there is only a 4 kΩ increase in the impedance despite the substantial delamination of graphene, which confirms the presence of Pt NPs even after longer ultrasonication (as shown in Fig. S3 in the supplementary notes).

## 4 Discussion

The multilayer graphene electrode surface is modified with Pt NPs to enhance its electrochemical characteristics. A spark ablation method is used to print NPs locally on the electrodes. This single-step process can be performed at room temperature in a dry environment. The surface modification practically enables the use of smaller electrodes with higher selectivity for neural recordings and allows the transfer of more charge via the electrode-tissue interface for neural stimulation under electrochemically safe conditions. Furthermore, the NP deposition at room temperature enables a stress-free NP coating deposition, which does not involve thermally introduced strain forces to the electrode^18^.

The highest NP surface density used in this work is 40% which still has an optical absorbance below 8%. The optical transmittance of graphene with the same growth condition was previously reported to be above 80%^11^. This confirms the potential use of graphene electrodes coated with NPs for future neuroscientific research such as optogenetics and optical imaging, as adding NPs on the graphene electrode surface is not expected to have a significant impact on the electrode’s optical transparency.

The results reported in this work demonstrate an improvement of the electrochemical characteristics of graphene electrodes by adding printed Pt NP coatings. The electrochemical performance is further improved for higher NP surface densities. This improvement is likely a result of the electrode surface area increase due to the presence of NPs and the fact that Pt NPs create a parallel conduction path in the electrode-electrolyte interface, overcoming the quantum capacitance of graphene^19^. CSC values are also known to be influenced by factors such as surface roughness, electrochemical surface area, and charge transfer mechanisms of the coating^37^. Previously reported monolayer graphene electrodes with electrodeposited Pt NP coatings have a similar area-normalized impedance for 15% NP density^19^. Unfortunately, in the aforementioned work, the stability of the coating on graphene was not investigated.

In this work, three different tests were used to assess the coating stability, electrochemically and mechanically. Continuous CV tests show that the electrodes with printed Pt NPs are electrochemically stable. Examination of the electrode surface after continuous CV measurements did not reveal any delamination or cracks on the graphene samples with NP coating. To further investigate the electrochemical stability of the electrodes, a larger number of CV cycles could be used.

Continuous VT tests do not suggest substantial delamination of the Pt NP coating from the graphene electrode surface. Similar impedance and CSC changes are observed for the graphene electrode without any coating which could be attributed to graphene delamination from the oxide layer. The reported impedance of the electrode with coating after 500,000 VT cycles is still lower than the impedance of graphene electrodes without any coating. This test should be repeated for the final implantable device as this device will not have a Mo layer underneath graphene which restricted this test due to Mo corrosion. The presence of the Mo layer probably had a negative impact on the results.

Ultrasonic treatment has been applied to the coated electrodes to assess the mechanical stability of the coating. Results from this test do not suggest NP delamination after the treatment, as indicated by optical inspection and impedance measurements. Surprisingly, graphene electrodes coated with NPs remain adherent to the underlying oxide layer, while graphene electrodes alone delaminate from it during the treatment. Ultrasonication has been used in literature to test the mechanical stability and adhesion of other coatings, such as PEDOT:PSS, on various electrode materials^39^. However, the ultrasonic vibration applied by this test to the electrodes is quite intense and harsh, and not representative to what an electrode will encounter in the body environment.

The test samples used for the stability assessment of the Pt NP coating on graphene in this work have not been optimised for these tests. In particular, the samples have been fabricated on a Si substrate where graphene sits on a silicon oxide layer, from which it delaminates during the continuous VT and ultrasonic treatments. This fact limited the intensity or duration of the treatment. In a practical application scenario of a neural interface, the oxide layer underneath the graphene electrodes is removed and substituted with parylene, as shown in^11^. Therefore, to ensure a more conclusive result the stability and adhesion tests should be repeated and extended for the final device. The delamination of the NP coating leads to the deterioration of the electrochemical characteristics over time, which consequently results in functionality loss. Moreover, the detached NPs may undergo biodispersion inside the body after implantation and can become toxic to the tissue. Additional treatments, such as prolonged immersion in a PBS solution, or dipping in an agarose gel^38^, could be added to the current suite, to further assess the long-term adhesion and stability of the coating for potential chronic applications.

If necessary, roughening the electrode surface prior to NP printing could be investigated in the future as a means to enhance the NP adhesion to the electrode surface. Previously, roughening the electrode surface of metal electrodes prior to PEDOT:PSS coating resulted in an increase in mechanical bonding between the electrode and its coating, thereby resulting in higher stability^39^.

The NP deposition technique presented in this paper yields a selective local modification of graphene neural electrodes. This opens up interesting possibilities when arrays of electrodes of various sizes are required during multimodal interaction with neural tissue. NP coatings come at the expense of less transparency, therefore could only be applied locally, only at e.g., very small electrodes, to enhance their recording performance, while larger electrodes on the same device can remain uncoated. Besides their effect on electrochemical characteristics, Pt NPs can be employed for local biosensing. This localization is not possible with electrodeposition techniques, where all electrodes on a device will be coated simultaneously. Besides, but crucially, the proposed technique is performed at room temperature and via a dry process, as a post-processing step. It is thus compatible with polymer substrates, an integral component of neural implants, as well as with a range of other processes and materials of the final device. These characteristics render this approach a unique tool for the enhancement of the performance of flexible neural implants.

## 5 Conclusions

To conclude, we present multilayer graphene neural electrode surface modification with Pt NPs using a spark ablation method. This method yields the local printing of NPs on an electrode surface without using a high temperature or wet processing. NP printing can be performed as a post-processing step to enhance the electrochemical characteristics of graphene electrodes further. The electrode showed 4.5 times lower impedance at 1 kHz after 40% NP coating on the surface (from 31.45 to 7.26 kΩ). The charge storage capacity (CSC), calculated based on a cyclic voltammetry (CV) test, was improved up to 15 times with 40% NP coating (from 233 *μ*C/cm^2^ to 3614 *μ*C/cm^2^). The maximum charge injection capacity (CIC), obtained by voltage transient (VT) measurements, also increased from 8.4 *μ*C/cm^2^ for graphene electrodes to 30.6 *μ*C/cm^2^ for graphene-coated electrodes with 40% surface density of NPs. NPs printed using this method yield electrochemical stability over 500 cycles of continuous CV measurements and 500,000 cycles of continuous VT tests. In addition, ultrasonic vibration of electrodes with NP coating shows better mechanical stability compared to graphene electrodes without any NPs. These results demonstrate selective NP deposition and local modification of electrochemical properties in neural electrodes for the first time, enabling the cohabitation of graphene electrodes with different electrochemical and optical characteristics on the same substrate.

## Acknowledgements

This research was supported by the POSITION-II project funded by the ECSEL JU under grant number Ecsel-783132-Position-II-2017-IA. The authors would like to thank the staff members of the Else Kooi Laboratory (EKL) for their support in the lab and VSPARTICLE B.V. for the use of their equipment.

