## Supplementary Notes for "Surface modification of multilayer graphene neural electrodes by local printing of platinum nanoparticles using spark ablation^†^"

#### Nanoparticle printing setting

Table S1. Nanoparticle printing setting

| Nominal NP surface density (%) | Calculated surface density (%) | Printing speed (mm/s) | Line width (μm) | Voltage (kV) | Current (mA) | Carrier gas flow (L/min) | Nozzle height (mm) | Nozzle diameter (mm) |
|---|---|---|---|---|---|---|---|---|
| 15 | 17.55 | 137.0 | 526 | 1 | 3 | $N_2$ (1.5) | 0.5 | 0.35 |
| 30 | 30.30 | 67.0 | 554 | | | | | |
| 40 | 39.90 | 38.3 | 637 | | | | | |

For the ease of comparison, 17.55% NP surface density was considered 15% in the text.

#### Conductivity measurement

Since the Pt NPs were printed as a line that goes from the insulating layer to the exposed electrode, the printed line may create a conducting path. To better understand this behaviour, a four-point probe measurement test was performed. A line pattern was printed over vertical metal tracks (100 nm gold with 10 nm chromium) on the samples. The measurement was performed by passing a current through two outer probes and measuring the voltage between two inner probes as shown in Figure S1 (a). The test was conducted twice for each printed line (with surface densities of 15, 30, 40, and 50%) at different locations. However, the distance between the inner probes was kept constant at 25 $\mu$m. The measurement was conducted from -500 mV to 500 mV with 8 mV steps.

Out of the eight measurement attempts (2 tests for each line sample), only 2 samples showed a current above the noise floor of the instrument. Hence, there is a conducting path along those measured areas. Two tests for the print sample of 50% NP show a conductive circuit. Presented in Figure S1 are the current-voltage curves measured for these closed circuits. Additionally, the 40% NP line did not have any current flowing on both tests, and the height difference between the exposed graphene electrode and the insulating layer (3 $\mu$m) on the final electrode samples likely severs any connecting path for print speeds of 40% NP and lower. Although, this is likely not the case for the 50% NP printed line with stable linear current-voltage characteristic curves. Therefore, the electrodes with 50% NP were removed from further characterization.

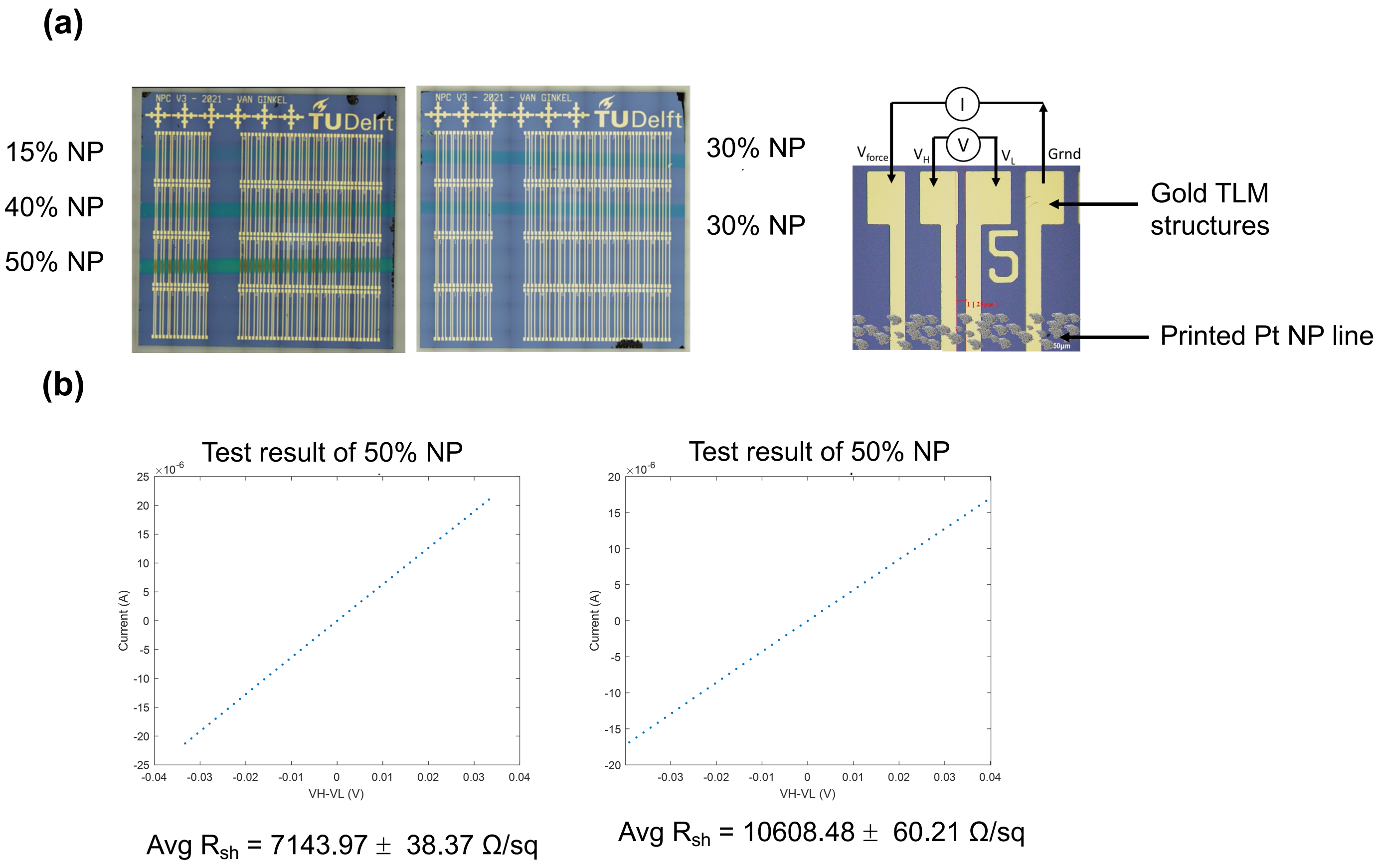


Figure S1. Conductivity measurement (a) Samples with Pt NP lines with the corresponding printing speed printed over gold TLM structures for the four-point probe measurement, (b) Results of the four-point probe measurements of 50% NP surface density printed lines.

#### Atomic force microscopy (AFM)

Table S2. RMS and mean surface roughness

| Nominal NP surface density (%) | RMS surface roughness (nm) | Mean surface roughness (nm) |
|---|---|---|
| 15 | 9.00 ± 0.87 | 7.11 ± 0.72 |
| 30 | 9.41 ± 1.37 | 6.96 ± 0.20 |
| 40 | 14.66 ± 7.31 | 12.03 ± 6.36 |

### Continuous CV test

Table S3. The results of continuous CV test for three electrodes per each group

| Electrodes | Z at 1 kHz (kΩ) Before | Z at 1 kHz (kΩ) After | Z change (%) | CSC (μC/cm²) | | | | | |
|---|---|---|---|---|---|---|---|---|---|
| | | | | Total | | | Cathodic | | |
| | | | | Before | After | Change (%) | Before | After | Change (%) |
| Graphene | 26.98 | 25.66 | -4.9 | 148 | 164 | 10.8 | 97 | 117 | 20.6 |
| | 34.60 | 33.61 | -2.8 | 99 | 101 | 2.0 | 48 | 56 | 16.7 |
| | 36.64 | 36.17 | -1.3 | 198 | 201 | 1.5 | 76 | 106 | 39.5 |
| Graphene + 15% Pt NPs | 17.19 | 19.83 | 15.4 | 738 | 675 | -8.5 | 596 | 548 | -8.0 |
| | 17.23 | 21.22 | 23.1 | 656 | 694 | 5.8 | 589 | 635 | 7.8 |
| | 18.03 | 17.99 | -0.3 | 767 | 834 | 8.7 | 646 | 703 | 8.8 |
| Graphene + 30% Pt NPs | 12.69 | 16.23 | 27.9 | 928 | 924 | -0.4 | 731 | 757 | 3.5 |
| | 10.90 | 13.33 | 22.3 | 797 | 789 | -1.0 | 678 | 659 | -2.8 |
| | 11.27 | 12.15 | 7.8 | 813 | 829 | 2.0 | 669 | 658 | -1.6 |
| Graphene + 40% Pt NPs | 7.87 | 8.62 | 9.5 | 879 | 1004 | 14.2 | 727 | 780 | 7.3 |
| | 7.62 | 8.53 | 12.0 | 952 | 1112 | 16.8 | 755 | 812 | 7.5 |
| | 9.22 | 8.80 | -4.6 | 917 | 1097 | 19.6 | 709 | 827 | 16.6 |

### EDX result

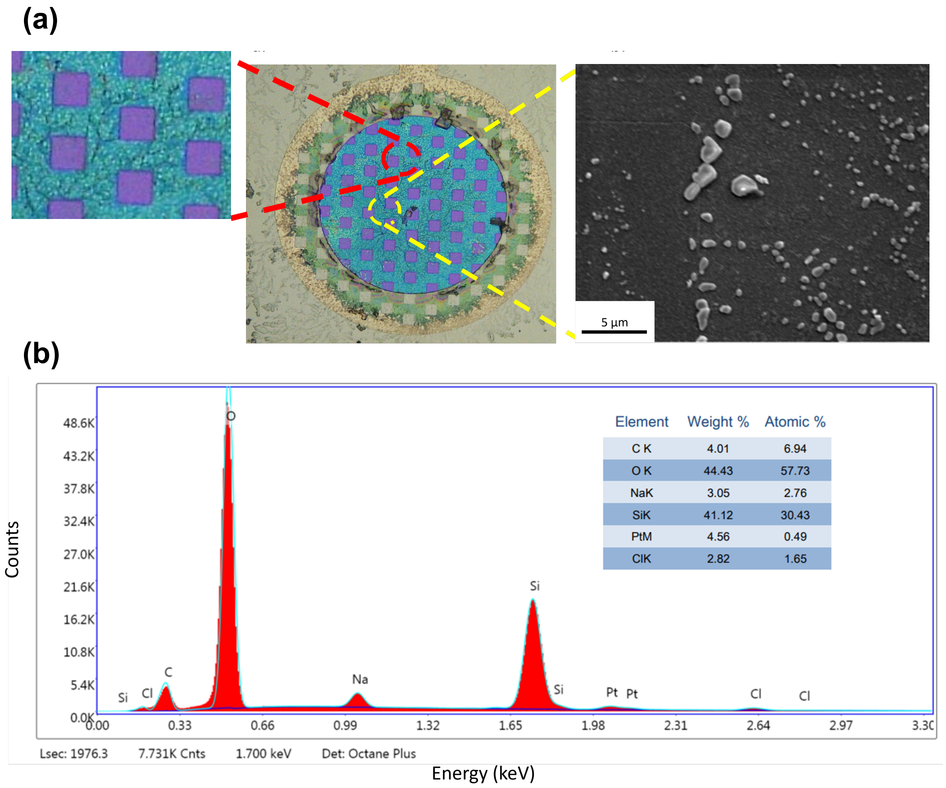


Figure S2. EDX result (a) Microscopy and SEM image of the electrode after the continuous CV (The zoomed-in optical image shows the dendritic pattern created by Na and Cl residues on the electrode surface), (b) EDX map spectrum and table of the present elements.

#### Ultrasonic test

The graphene electrodes without NPs showed partial to complete delamination of the layer after only 2 minutes of the ultrasonic test as shown in Figure S3 (a). This is due to the poor adhesion of graphene to the underlying oxide. Samples with NP coatings were able to sustain ultrasonication longer than samples without NPs.

After 5 minutes of ultrasonication, there was visible delamination of the graphene layer on samples with NPs. However, the impedance (at 1 kHz) of the electrode (as shown in Figure S3. (b)) is 13 kΩ which is still much lower than the average impedance of the graphene (34 kΩ ), thus implying that there are still Pt NPs on the electrode surface. However, the Pt NP surface density after this duration is unknown. The impedance value is still low (26.25 kΩ) after the 7 minutes of ultrasonication and increases (38.7 kΩ) after 10 minutes. Despite the substantial delamination of the graphene layer, the impedance only increased by about 4 kΩ. This suggests that there may still be some Pt NPs on what is left of the graphene layer. There was also a color change on the graphene after 12 minutes, possibly caused by substantial delamination of the Pt NPs.

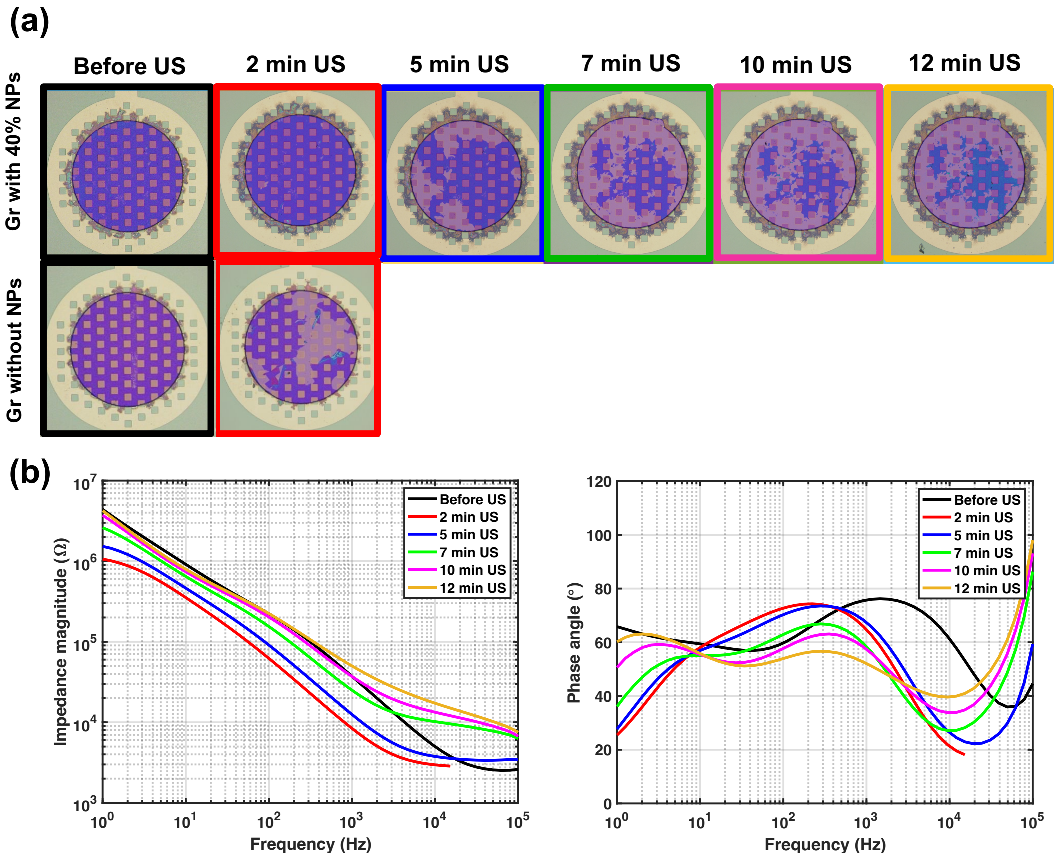


Figure S3. Ultrasonic test (a) optical image of graphene electrodes with and without NPs before and after 12 minutes of ultrasonication, (b) Impedance magnitude and phase plot of graphene electrodes with 40% NPs before and after 12 minutes of ultrasonication.
